# Gap scheduling of a PARP inhibitor and nanoparticle TOP1 agent combination avoids synergistic bone marrow toxicity

**DOI:** 10.1101/2025.04.30.651268

**Authors:** Lenka Oplustil O’Connor, Anderson Wang, Claire Sadler, Jennifer Barnes, Rajesh Odedra, Aaron Smith, Gareth Hughes, Alan Lau, Andres Tellez, Scott Eliasof, Elaine Cadogan, Mark J. O’Connor

## Abstract

Although combinations of DNA damage response inhibitors (DDRi) and DNA damaging chemotherapy enhance cytotoxicity in cell-based systems, clinical success has been limited by overlapping bone marrow toxicities. Here, we show that a tumor-targeted nanoparticle camptothecin CRLX101, administered concurrently with DDRi, enhances anti-tumour efficacy but also increases bone marrow toxicity in preclinical models. Using rat bone marrow progenitor cells as biomarkers and leveraging differential repair kinetics of CRLX101-induced DNA damage in tumour and bone marrow, we identified a gap schedule of the PARP inhibitor olaparib and CRLX101 that enhanced efficacy over single agents but demonstrated a reduced combination marrow toxicity. A clinical trial has been designed using the gap schedule identified here and represents a template that can be used to successfully deliver DDRi with tumor-targeted chemotherapy in combination.

## Introduction

DNA damaging chemotherapy has been at the centre of cancer therapy for over seven decades (Cheung-Ong et al., 2013; DeVita and Chu, 2008). Underlining the preferential sensitivity of tumours compared to normal tissues are deficiencies in the DNA damage response (DDR), a collective term for the intra- and inter-cellular signaling events involved in the detection and repair of DNA damage (Ciccia and Elledge, 2010; Jackson and Bartek, 2009). In spite of the cancer-specific DDR deficiencies (Jackson and Bartek, 2009), the levels of exogenously generated DNA damage caused by systemic chemotherapies still results in significant and unwanted side effects. For this reason, there has been an attempt over the last twenty years or so, to identify specific inhibitors of the DDR in order to exploit the same cancer-specific DDR deficiencies (Curtin, 2012; O’Connor, 2015). In the case of inhibitors of poly (ADP-ribose) polymerase (PARP), this has led to the first approved DDR-based medicines, where PARP inhibitors (PARPi) have demonstrated monotherapy activity (Coleman et al., 2017; de Bono et al., 2020; Golan et al., 2019; Ledermann et al., 2014; Litton et al., 2018; Mirza et al., 2016; Robson et al., 2017) through a mechanism described as synthetic lethality (Fong et al., 2009; Lord and Ashworth, 2017). However, the potential for PARP inhibitors goes well beyond monotherapy in HRR-deficient cancers and includes the potentiation of DNA damaging chemotherapy in a broader range of tumours (Pilie et al., 2019). Moreover, in addition to inhibitors of PARP, there are now several other DDR targeted agents being developed in the clinic (O’Connor, 2015), including those agents that inhibit WEE1 (Hirai et al., 2009), ATR (Min et al., 2017; Reaper et al., 2011; Wengner et al., 2020), CHK1 (Angius et al., 2020), ATM (Durant et al., 2018; Riches et al., 2020), DNA-PK (Fok et al., 2019; Wise et al., 2019) and Aurora Kinase B (Ashton et al., 2016).

While there are no shortages of preclinical examples where DDR inhibitors (DDRi) enhance the anti-tumour activity of chemotherapies, in most cases, poor tolerability for these combinations is underestimated because of the use of sub-optimal preclinical models. For example, mouse tumour efficacy models will not accurately reflect the potential impact on bone marrow since mice preferentially use different (more efficient but error-prone) DDR pathways in bone marrow stem cells than either rats or humans (Lane and Scadden, 2010; O’Connor et al., 2016). For this reason, rats represent a better preclinical safety model than mice to assess bone marrow toxicity. When DDRi are combined with chemotherapy, the experience from the clinic is that the overlapping toxicities with DDRi present a significant challenge and currently there have been few successful trials demonstrating efficacy coupled with acceptable tolerability. This point is highlighted by the fact that currently there isn’t a single approval for a targeted DDR agent in combination with chemotherapy.

One approach to improve the therapeutic window of chemotherapy is to target the DNA damaging agent to the tumour and/or avoid the bone marrow compartment. Approaches to achieve this currently in development include re-formulation (e.g. through liposomal or nanoparticle approaches (Wang et al., 2012)) or conjugation of the chemotherapy war-head to an antibody that preferentially targets a tumour surface marker (antibody-drug-conjugates or ADCs; (Beck et al., 2017; Thomas et al., 2016)).

Here, we investigated the combination of a nano-particle camptothecin, CRLX101 (Weiss et al., 2013), with various DDRi, including the PARPi olaparib (Menear et al., 2008), WEE1i adavosertib/AZD1775 (Hirai et al., 2009), ATRi ceralasertib/AZD6738 (Foote et al., 2018), ATMi AZD0156 (Riches et al., 2020), DNA-PKi AZ’6119, a precursor of AZD7648 (Fok et al., 2019), and Aurora kinase B inhibitor AZD1152 (Wilkinson et al., 2007). We chose CRLX101 for combination studies based on its monotherapy activity, preferential accumulation in tumors, and favorable activity compared to conventional topoisomerase I inhibitors (TOP1i) (Clark et al., 2016; Pham et al., 2015; Weiss et al., 2013). Triaging the combinations *in vitro,* we identified the ones demonstrating increased efficacy. We then assessed the most promising combinations in a rat bone marrow safety model to identify the best tolerated combination of CRLX101 and DDRi. Optimization of the combination dose and schedule were carried out using rat bone marrow and mouse small xenograft tumour models which identified a starting point for clinical assessment. A phase 1 study has been designed to assess the predictive value of our preclinical approach.

## Results

### DDR inhibitors are differentiated in both the degree and mechanism of CRLX101 potentiation

CRLX101 is a targeted topoisomerase I inhibitor (TOP1i) in clinical development based on a nanoparticle camptothecin formulation (Clark et al., 2016). Small cell lung cancer (SCLC) is a tumour type of interest for CRLX101 development as these tumors are responsive to TOP1 inhibitors, which are part of standard therapeutics topotecan. Several. SCLC also represents a tumour type of high unmet clinical need. To evaluate the benefit of combining CRLX101 with different DDR inhibitors, we chose to use the SCLC cell line model NCI-H417, because the response to single agent CRLX101 had previously been characterized and importantly, the NCI-H417 cell line could be used to generate an in vivo xenograft model.

We treated the NCI-H417 cells with CRLX101 in combination with a PARP inhibitor (olaparib), a WEE1 inhibitor (AZD1775), an ATM inhibitor (AZD0156), an ATR inhibitor (AZD6738), a DNA-PK inhibitor (AZ’6119) and an Aurora kinase B inhibitor (AZD1152). Cells were exposed to CRLX101 for 48h, followed by drug wash out. A fixed dose of DDR inhibitor (shown to provide greater than 95% target inhibition) was added concurrently to varying concentrations of CRLX101 on day 1 and replenished after 48h. Cell viability was determined at day 7 using an MTT assay (Figure 1A). The PARPi, ATRi and ATMi all significantly potentiated the activity of CRLX101, resulting in a decrease in the GI_50_ of CRLX101 by 3.2-fold, 3.6-fold and 7.8-fold respectively (Figure 1A). A significant proportion of cells (∼20-30%) were not killed by CRLX101 treatment alone, even at the higher concentrations. Combinations with the PARPi, ATMi, or ATRi shifted the GI_50_ curve but did not result in any increase in this maximum cell kill value. Combination with the WEE1 inhibitor (AZD1775), although not significantly shifting the GI_50_ curve, did result in a more potent cell kill (>90% cells). No obvious combination benefit was seen with DNA-PK inhibitor (AZ’6119) or the Aurora B kinase inhibitor (AZD1152). Combining a fixed dose of CRLX101 (3nM) with varying concentrations of olaparib or AZD1775 similarly demonstrated a shift in the efficacy curves of WEE1i and the PARPi. This effect was particularly striking for the olaparib dose response combination, where 0.3µM concentration of olaparib was sufficient to achieve greater than 50% growth inhibition (Supplementary Figure S1) suggesting that lower doses of olaparib could be used *in vivo* or in the clinical setting, in case there is a necessity to dose reduce due to normal tissue toxicities. Olaparib was also seen to lower the GI_50_ of CRLX101 by 2.6-fold in another cell line model, the ovarian cancer model POE4 (Supplemental Fig. S1).

**Figure 1:**
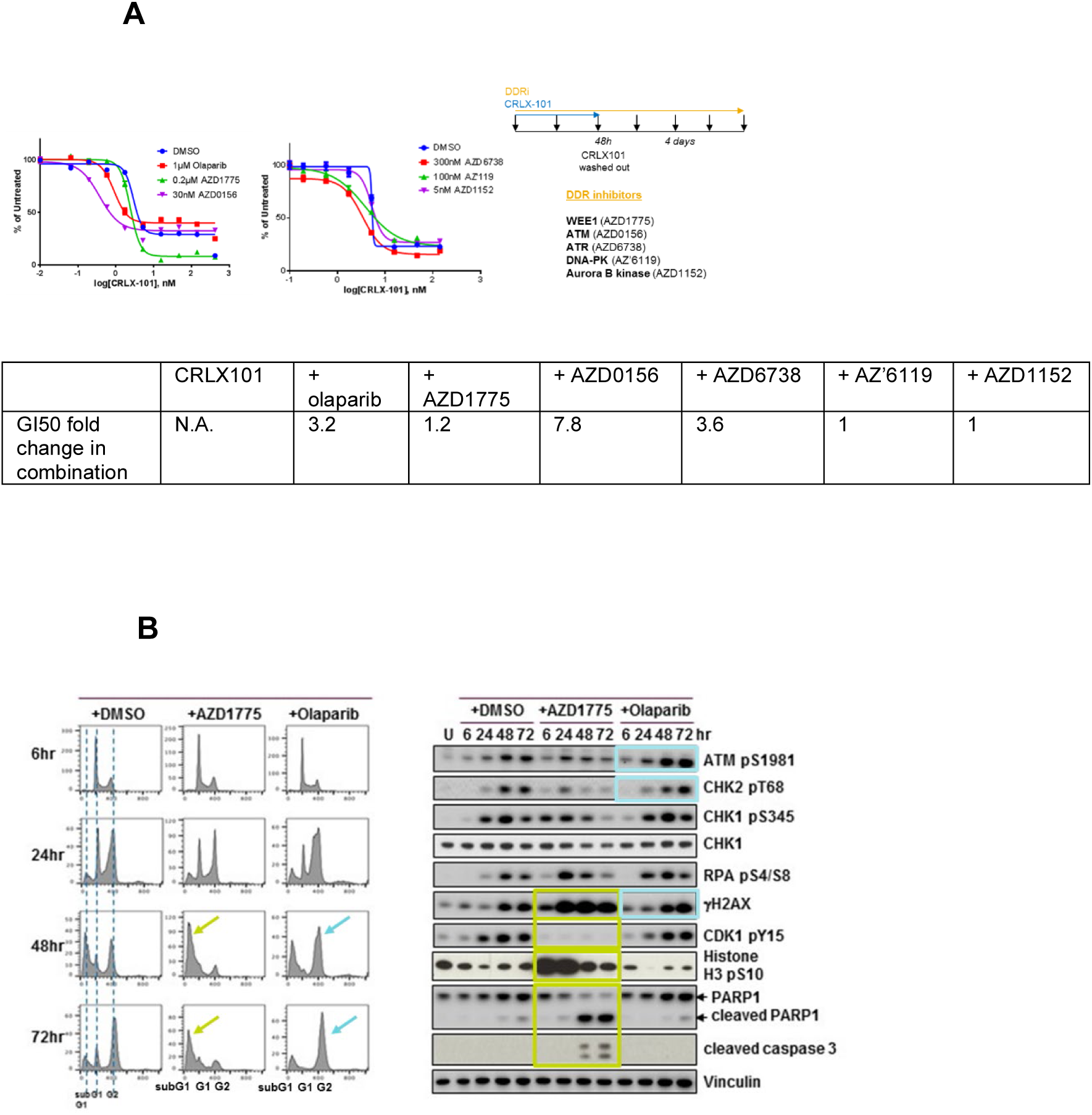
**(A) Potentiation of CRLX101 by different DDR inhibitors.** SCLC cell line NCI-H417a treated with CRLX101 for 48h before washing out the drug. DDR inhibitors were added concurrently with CRX101 and replenished after 48h. Growth inhibition (GI_50_ values) were determined at day 7 using the MTT assay. **(B) Olaparib and AZD1775 potentiate the efficacy of CRLX101 through different mechanisms.** NCI-H417a cells were treated with CRLX101 for 48 hours either alone or in combination with indicated DDR inhibitors. Following washout of CRLX101, DMSO or DDR inhibitors were replenished. Samples were harvested at indicated timepoints for cell cycle status analysis by flow cytometry (left panel) or DDR biomarker activation analysis by immunoblotting (right panel).

To understand the mechanism of action of different DDRi when combined with CRLX101, we initially compared combination treatments for CRLX101 with PARPi or WEE1i. First, we harvested NCI-H417a cells at various timepoints following treatment and analyzed cell cycle changes (Figure 1B, left) as well as the activation of DDR biomarkers (Figure 1B, right). CRLX101 activated a number of DDR markers, including pATM, pCHK2, pCHK1, pRPA and γH2AX. The treatment of CRLX101 (+DMSO control) resulted in a transient decrease in phospho-Histone H3 (pHH3), likely reflecting the induction of DNA damage (γH2AX) and replication stress (pS4/8 RPA) in S-phase. By contrast, the combination of CRLX101 with the WEE1 inhibitor AZD1775 leads to an initial increase in cells entering mitosis, since WEE1 inhibition, while increasing DNA damage and replication stress (γH2AX, pRPA and pCHK1), also overrides the G2/M checkpoint as indicated by pCDK1 inhibition and previously described (Young et al., 2019). In this study, following WEE1i treatment, an initial early mitotic entry of cancer cells that have already undergone DNA replication and were in late S or G2 phase occurred, followed by a reduction in mitotic entry as cells that had not undergone replication arrested in early S phase. In the NCI-H427a cells combination of CRLX101 with WEE1 inhibition therefore resulted in aberrant mitotic entry and enhanced cell death. The latter is indicated by increases in cleaved PARP and cleaved caspase 3 levels and extensive sub-G1 peak (Figure 1B). By contrast, the olaparib combination enhanced CRLX101-induced DNA damage, as indicated by elevated pATM and γH2AX, as well as activating the G2/M checkpoint (pCHK1 and pCDK1), with a concomitant reduction in pHH3. Combining PARP inhibitor with CRLX101 thus likely results in more DNA double-strand breaks and a prolonged cell cycle arrest in the late S and G2 phases (Figure 1B).

### Increased hematological impact of concurrent CRXL101 and olaparib combination schedule

The strong mechanistic rationale for combining TOP1i with olaparib (Murai et al., 2014; Pommier et al., 2016) has previously encouraged the initiation of several Phase 1 clinical studies to investigate the tolerability of olaparib in combination with irinotecan (Chen et al., 2016) or topotecan (Samol et al., 2012). However, these combinations were not developed any further in the clinic, due to dose-limiting hematological adverse effects and a resulting sub-therapeutic combination maximum tolerated dose (MTD).

The pharmacokinetics and exposure levels of CRLX101 in both mouse and rats have previously been published and compared to exposures at different drug dose levels in humans that identified the human MTD (15mg/m^2^) equivalent dose for both species, namely 5mg/kg in the mouse and 2.5mg/kg in the rat. To assess CRLX-101 induced bone marrow toxicity in rats we used either 2.5mg/kg (MTD), 2mg/kg (80% MTD) or 1.25mg/kg (50% MTD), while for mouse xenograft efficacy studies, we used either 5mg/kg (MTD) or 4mg/kg (80% MTD).

Assessment of bone marrow toxicity in immunocompetent RccHan:WIST rats using CRLX101 and olaparib as single agents, and compared to topotecan treatment, is shown in Figure 2A. There is little effect of either olaparib or CRLX-101 at 48h. By comparison, topotecan treatment at day 2 can be seen to result in a clear reduction in the myeloid progenitor (CD45^+^) population. The nadir for the effect on these myeloid progenitors peaked at Day 6, and this is also shown in Figure 2A. However, the combination of olaparib and CRLX-101 did demonstrate much greater bone marrow toxicity, compared to either single agent alone. The nadir effect for the olaparib CRLX-101 combination toxicity occurred at day 7 and the effects at this time point are shown in Figure 2B. Compared to CRLX-101 treatment alone, the MTD and 50% MTD of CRLX-101 when combined with olaparib The combination effect of olaparib and CRLX101reduction in combination in the bone marrow, involved using immunocompetent RccHan:WIST rats, treated with 2mg/kg qdx1 single agent CRLX101 or olaparib at 100mg/kg qdx7, or with concurrent combination of the two agents (Figure 2).

**Figure 2:**
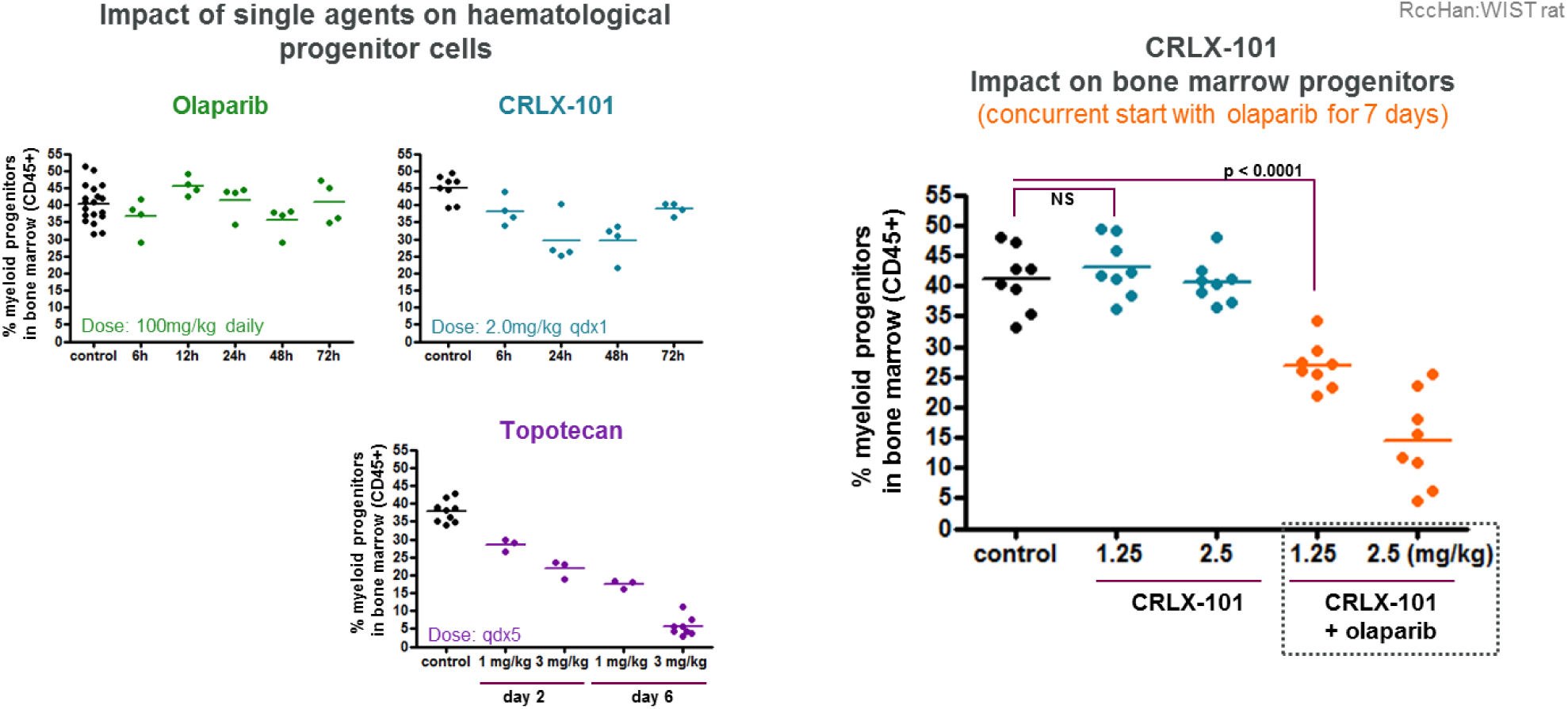
Increased hematological impact of CRLX101 in concurrent combination with olaparib. RccHan:WIST rats were treated with individual agents or simultaneously with CRLX101 and olaparib. Bone marrow cells were harvested from the rat femur and analyzed for frequencies of nucleated-myeloid LDS-751^pos^ CD45^pos^ progenitor cells by flow cytometry.

Bone marrow cells were harvested on day 7 from a femur and analyzed for frequencies of nucleated-myeloid (LDS-751 positive/ CD45 positive) progenitor cells by flow cytometry. We also included a topotecan-treated group (1mg/kg and 3mg/kg qdx5) as a reference, clinically relevant, standard-of-care agent and schedule. Both CRLX101 and olaparib were well tolerated as single agents. However, concurrent combination resulted in a dose-dependent decrease in bone marrow progenitors in the rat bone marrow model using the preclinical equivalent MTD of 2.5mg/kg and a half equivalent MTD of 1.25mg/kg. Topotecan, even as a monotherapy, led to a pronounced loss of bone marrow cells over the treatment period (Figure 2). The effect of the concurrent combination of olaparib with CRLX101 was also observed in peripheral blood (Supplementary Figure S2). Assessment of the hematological effects of two other DDR inhibitors, namely the WEE1i (AZD1775) and the ATMi (AZD0156), was also carried out. WEE1i was chosen, since this gave the greatest maximum cell kill in combination with CRLX101 of all the DDRi tested, while ATMi was chosen because this gave the greatest level of potentiation (nearly 8-fold). Combinations of both agents demonstrated a greater degree of effect on the bone marrow than olaparib (Supplementary Figure S2).

### CRXL101 treatment generates different DNA damage induction profiles in bone marrow vs. tumour

Our initial data from the rat bone marrow response studies (Fig. 2) indicated a potential risk that concurrent combination of CRLX101 with olaparib would lead to exacerbated hematological toxicity. Therefore, we sought to determine whether rational sequencing of the two agents could be explored to improve bone marrow tolerability, while not compromising on anti-tumour efficacy. To evaluate the effect of CRLX101 on DNA damage in bone marrow versus tumour cells, we treated preclinical models - rats for bone marrow analysis, and NCI-H417 tumour bearing nude mice – with equivalent, clinically relevant exposures of CRLX101 and harvested bone marrow and tumour tissue between 6h – 72h post dosing. The samples were processed for immunohistochemistry staining and labeled with antibodies against γH2AX (indicative of DNA double-strand breaks) or cleaved caspase 3 (indicative of apoptosis). Plasma, tumour and bone marrow cells were collected for pharmacokinetic analyses by mass spectrometry (Supplementary Figures. S3 and S4). We found that only a small fraction of CRLX101 (<1% plasma levels at time points 6-72h) penetrates the bone marrow, resulting in only low levels of transient DNA damage (indicated by γH2AX from the 6h to 24h timepoints) with little observed loss of bone marrow cellularity (Figure 3A). In contrast, CRLX101 is retained for a much longer time in tumour cells, leading to sustained DNA damage and associated cell death (Figure 3A).

**Figure 3:**
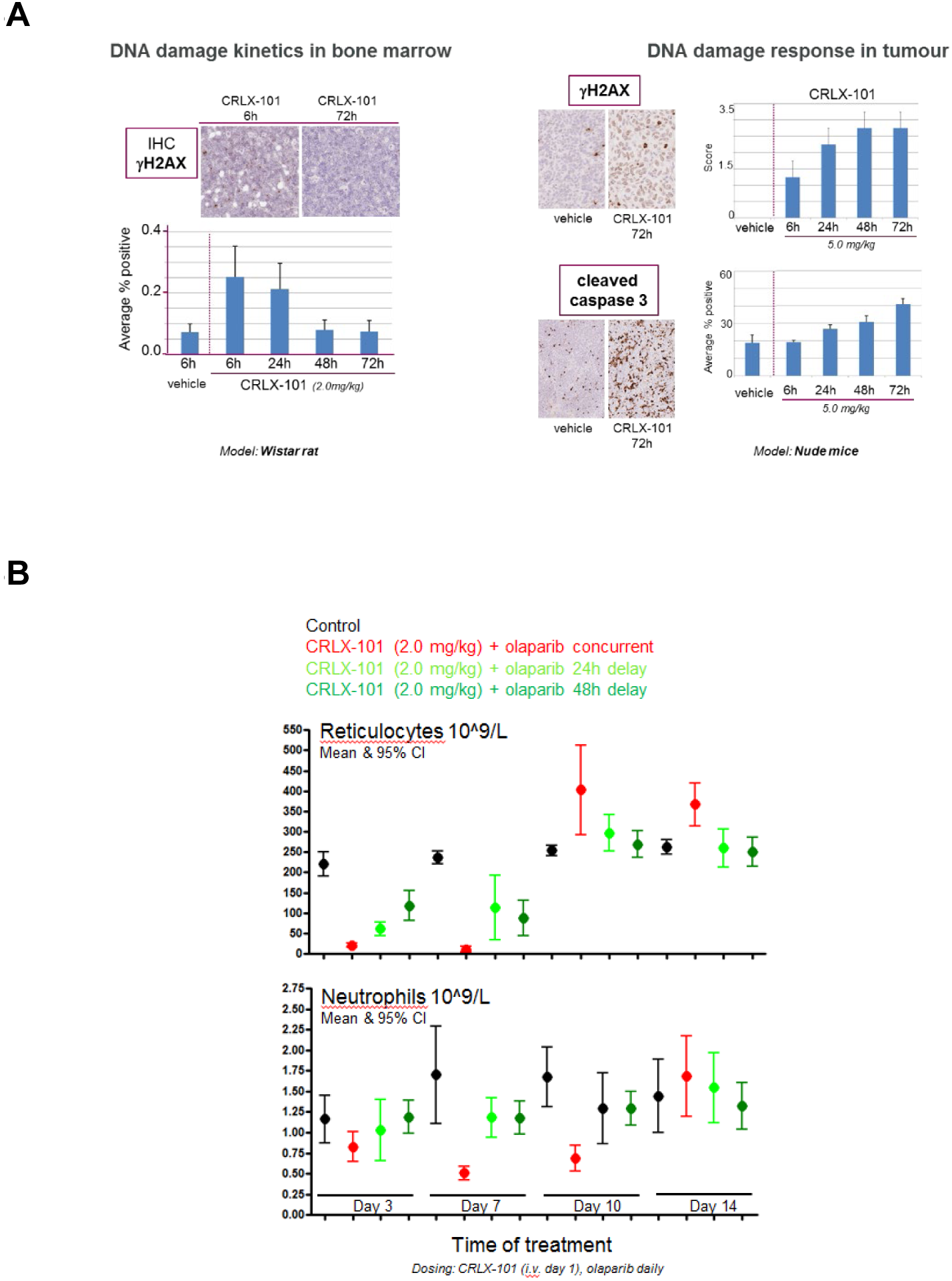
**(A) CRLX101 exposure leads to a different profile of DNA damage induction in bone marrow cells vs tumour.** Animals received a clinically relevant dose of CRLX101 and the tissue (bone marrow from WT rats, tumour from NCI-H417a bearing nude mice) harvested between 6h – 72h. The samples were processed for immunohistochemistry staining and labeled with antibodies against γH2AX (DNA double-strand breaks) or cleaved caspase 3 (apoptosis). **(B) Sequenced (gap) schedules of CRLX101 and olaparib decrease the impact on bone marrow.** Wistar rats were treated with a single dose of CRLX101 and either simultaneously or with 24h and 48h delay before daily treatment with olaparib. Peripheral blood was collected from tail vein and analysis was performed on the Siemens Advia 2120i haematology analyser.

### A sequenced (gap) schedule of CRLX101 with olaparib reduces the impact on bone marrow

The pharmacodynamic biomarker analysis outlined in Figure 3A demonstrated clear differences following a single dose of CRLX101, between DNA damage induction and recovery in bone marrow vs. tumour tissue. (Figure 3A). Based on these results, we hypothesized that dosing olaparib with at least a 24h gap following CRXL101 treatment could allow bone marrow cells to recover from the initial CRLX101-induced damage before olaparib had the chance to potentiate the effect, thus reducing the chance of cell death and facilitating hematological recovery. Given the dose-dependent effects seen in Figure 2, we chose an intermediate dose of CRLX101 for the gap schedule of 2mg/kg, or 80% of the equivalent clinical MTD. While 24h or 48h following CRLX101 treatment should provide the potential for bone marrow DNA damage resolution, the γH2AX biomarker data indicated this would not be the case for tumour cells since sustained and elevated levels of DNA damage, as well as the continued presence of the compound, were detected at 24h and at later time points. Thus, it would be predicted that even with a 24h or 48h gap, the effects of CRLX101 could be further potentiated by olaparib. To test this hypothesis, we treated rats with a single dose of CRLX101 and then dosed olaparib simultaneously or with a 24h or 48h delay. Peripheral blood was collected from tail veins and blood differentials analysed. Combination using a delayed or gap schedule of 24h or greater between the CRLX101 and the olaparib had a sparing effect on peripheral blood cells, in terms of both the extent of the nadir and the time to recovery (Figure 3B).

### A sequenced (gap) schedule of CRLX101 with olaparib maintains anti-tumour efficacy

A previous SCLC in vivo study of concurrent treatment of olaparib with CRLX101 in the xenograft model NCI-H1048 demonstrated that the combination anti-tumour activity was more effective than CRLX101 treatment alone, with 0/7 tumour progressions in the combination cohort versus 7/7 for CRLX101 as a single agent (Supplementary Figure S5). To test whether gap scheduling of CRLX101 and olaparib can maintain enhanced anti-tumour efficacy of the combination over single agent CRLX101 alone, we tested the response *in vivo* in a small cell lung cancer NCI-H417 subcutaneously implanted xenograft model using 4mg/kg CRLX101 (one dose/week), which is 80% of the clinical MTD equivalent in mice, with daily dosing of 100mg/kg olaparib (MTD equivalent) following a 24h gap for two weeks. As a comparator, we used the 4mg/kg single agent dose of CRLX101, single agent olaparib, and topotecan at the MTD for this agent of 1mg/kg for five days, both as a single agent and in combination with olaparib in a 24h gap schedule. Individual tumour volumes were measured regularly up to 83 days. This study demonstrated that the efficacy of CRLX101 in combination with olaparib is better compared to CRLX101 alone and is significantly more efficacious than MTD topotecan treatment (Figure 4A).

**Figure 4:**
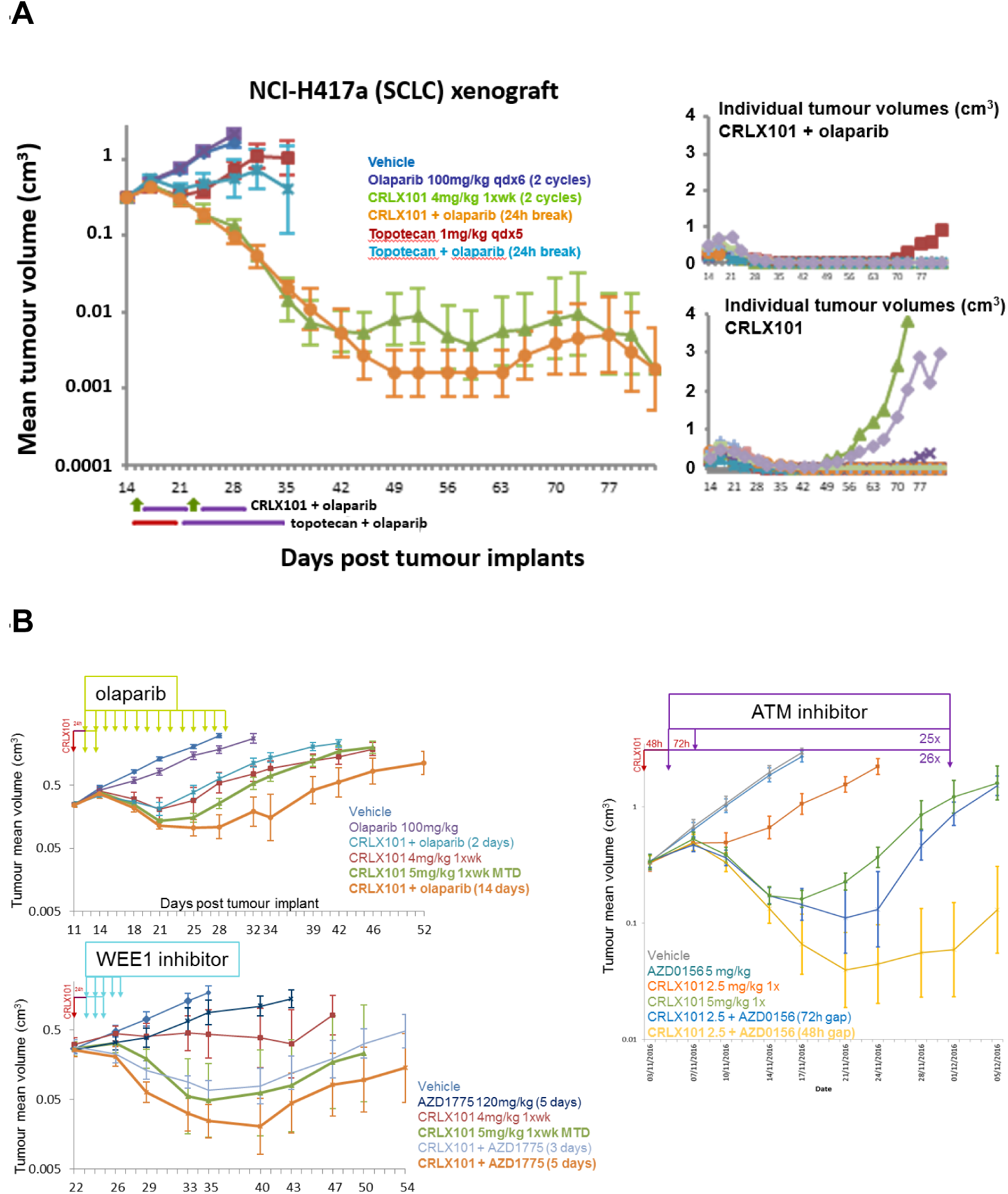
**(A) Gap scheduling of olaparib following CRLX101 treatment maintains anti-tumour efficacy compared to single agent CRLX101 alone.** Human small cell lung cancer cells (NCI-H417a) were implanted subcutaneously in athymic Fox1-nu mice. Animals were dosed with individual agents or combination of CRLX-101 (1xwk) + olaparib (qdx10, 24h after) or topotecan (qdx5) + olaparib (qdx14, 24h after). Individual tumour volumes were measured regularly up to 83 days. **(B) Extending the duration of olaparib treatment following CRLX101 treatment maximises the anti-tumour effect of the combination.** Mice implanted with NCI-H417a tumour cells were dosed with single dose of CRLX101 + olaparib (100 m/kg) for 2 days or 14 days (top panel) or CRLX101 + AZD1775 (120mg/kg) for 3 or 5 days (bottom panel) starting 24h after the CRLX101 or CRXL101 + AZD0156 (5mg/kg) for up to 26 days.

Given that the *in vivo* model we were using did not have an identified homologous recombination repair (HRR) deficiency that would provide olaparib single-agent sensitivity, we asked the question of whether there would be any benefit in extending the dosing of olaparib beyond a couple of days after CRLX101 treatment. Mice implanted with NCI-H417a tumour cells were therefore treated with a single dose of CRLX101 (4mg/kg) and after 24h, olaparib (100mg/kg) for either 2 days or 14 days. The data depicted in Figure 4B clearly demonstrate that dosing olaparib for 14 days is significantly better than for just 2 days, even in a homologous recombination repair-proficient background. The most likely explanation for the enhanced potentiation provided by extended olaparib treatment is that CRLX101 is retained in tumours and continues to generate DNA damage over time, the effects of which are enhanced by the continued presence of olaparib.

Finally, to extend our *in vitro* data suggesting that the WEE1i and ATMi may also potentiate the activity of CRLX101, we further tested these two combinations in the NCI-H417a tumour xenograft model (Figure 4B). Mice were treated with CRLX101 (4mg/kg) plus AZD1775 (120mg/kg) for 3 or 5 days starting 24h after the CRLX101. The combination of reduced dose of CRLX101 (4mg/kg) with 5 days of AZD1775 dosing led to better efficacy than the 3-day schedule and was also more potent than CRLX101 monotherapy at its maximum tolerated dose (5mg/kg). However, clinical data on adavosertib/AZD1775 suggests it will not be possible to be able to deliver the 120mg/kg equivalent dose in humans for five days (Do et al., 2015). For the ATMi, initial tolerability studies in rats showed that a combination with CRLX101 dosed 24h apart was not tolerated. Reflecting on this observation and the fact that the ATMi was the most potent potentiator of CRLX101 activity *in vitro* (Figure 1), we focused on 48h and 72h gap schedules and further reduced the dose of CRLX101 to 50% MTD equivalent (2.5mg/kg in mice) for this combination. It was encouraging to see that the combination of 2.5mg/kg of CRLX101 with AZD0156 dosed 48h apart led to tumour regression, significantly greater than full dose of CRXL101 alone (Figure 4B). However, bone marrow tolerability would be a potential concern. The anti-tumour efficacy was still maintained with the 72h gap schedule; however, the effect was just marginally greater that CRLX101 alone when dosed in its MTD.

Taken together, our data suggested that while there were a number of options for DDRi combinations with CRLX101, the most likely to succeed clinically was the PARPi olaparib combination. The preclinical studies suggested a combination dose and schedule of 80% of the clinical MTD equivalent of CRLX101 followed by a 48h gap and then daily olaparib using a dose escalation approach to get as close to the MTD as possible until two days before the next CRLX101 treatment begins, to avoid concurrent dosing at the beginning of the second cycle. Based on these insights from preclinical studies, a phase I clinical trial combining olaparib and CRLX101 has been designed and carried out to assess the gap schedule approach (NCT02769962).

## DISCUSSION

Many standard-of-care (SoC) chemotherapies act by generating DNA damage that has the potential to be enhanced by inhibitors of the DDR. However, overlapping toxicities, in particular bone marrow toxicity, has hampered the ability to combine these agents. Attempting to identify an effective and sufficiently well tolerated dose and schedule in the clinic can take many years and still fail to identify a combination that is clearly better than the full dose SoC chemotherapy alone. A good illustration of this is the combination of the PARPi olaparib with carboplatin and paclitaxel, where a Phase I trial (NCT00516724) began in 2007, took many years to complete and involved 189 patients. Only recently has this trial been reported (van der Noll et al., 2020) and the eventual conclusion was that olaparib in combination with carboplatin and/or paclitaxel resulted in increased hematologic toxicities, making it challenging to establish a dosing regimen that could be tolerated for multiple cycles without dose modifications. Another example, where there is a clear mechanistic rational for combination but where enhanced bone marrow toxicity has meant no optimal recommended Phase II dose has been identified that could be taken forward, is the combination of PARPi with TOP1i such as irinotecan and topotecan (Murai et al., 2014). Several clinical trials (Chen et al., 2016; Dhawan et al., 2017; Hendrickson et al., 2018; Kummar et al., 2011; LoRusso et al., 2016; Rajan et al., 2012; Samol et al., 2012) have tried and failed to combine PARPi and TOP1i at doses close to their individual MTDs with dose limiting myelosuppression precluding administration of >20% of a PARPi MTD dose.

The initial preclinical rat bone marrow studies (Figure 2) indicated that even with the nanoparticle targeting of the TOP1i, concurrent treatment was having a synergistic toxicity effect, greater than either agent alone. However, by deploying a gap schedule, 80% of the MTD equivalent of CRLX101 could be tolerated in combination with an MTD equivalent of olaparib, which has led to the initiation of a clinical trial to test this gap schedule combination (Clinical trials.gov reference NCT02769962, where CRLX101 was renamed EP0057).

The gap scheduling approach exemplified here also has the potential to work for PARPi with other targeted DNA damaging agents and for other DDRi. For example, one of the strongest potentiators of TOP1i is ATM inhibition, and we demonstrate here for CRLX101, potentiation both *in vitro* (Figure 1) and *in vivo* (Figure 4). However, the flip side to this significant potentiation is greater combination bone marrow and peripheral haematological toxicity. No doubt ATM inhibitors should be assessed with effective targeted delivery of DNA damaging agents, but we anticipate a gap scheduling approach is also likely to be required in this scenario. Moreover, nanoparticle technologies that significantly enhance the systemic half-life of their chemotherapy payload are clearly going to be difficult to use with the gap scheduling approach for DDRi combinations outlined here, which very much depends upon the differentiation between tumor and normal tissue levels of DNA damage and repair kinetics. In addition to nanoparticle conjugated chemotherapies, such as the CRLX101 used in this study, other technologies that can deliver DNA damaging chemotherapy specifically to tumor, such as antibody drug conjugates (ADCs), could benefit from gap combination schedules with DDR inhibitors. In addition to tumour specific receptors another important requisite will be both the half-life of the ADC as well as the stability of the ADC linker in order to avoid continuous exposure of bone marrow and other normal tissues to the ADC payload.

Finally, one additional aspect of DDRi combinations that is likely to benefit from gap or non-overlapping schedules is the DDRi-DDRi combination. There are an increasing number of examples where combining two DDR inhibitors is proving effective in exploiting tumor DDR-deficiencies. For example, PARPi, WEE1i and ATRi combinations have been shown to be effective pre-clinically in several tumour settings (Fang et al., 2019; Lallo et al., 2018; Young et al., 2019). A number of these combinations are currently being tested in the clinic. However, concurrent scheduling is also limiting the extent of single agent dosing that can be given due to enhanced bone marrow toxicity. An alternative approach based on differences between DDRi effects in tumour vs normal tissue have recently been described (Fang et al., 2019), where effects of DDR agents were seen to last for several days in tumours but not haematological cells, leading to greater dependency on the second DDR agent specifically in the tumour and opening the way for alternating scheduling of DDRi to be assessed.

In summary, DDR inhibitor combinations, either with DNA damaging agents or other DDR inhibitors, have significant therapeutic potential but are nevertheless challenging. The work presented here has highlighted how an assessment of preclinical models of both tumour and normal tissue responses, such as those of rat bone marrow, can provide dose and scheduling insights that can significantly enable combination testing in the clinic.

## Acknowledgements

We would like to thank those current and past AstraZeneca scientists, who are not co-authors but who nevertheless contributed towards the work, namely Jamie Reens, Catherine Wilkinson, Anna Cronin, Richard Knights and Pete Newham.

## Author Contributions

Original conceptualization and methodology for the preclinical studies was provided by MOC and LOC. Experimental investigation was provided by LOC, AW, CS, JB, RO, AS, GH, JR, AL and EC, with overall supervision from MOC.

## METHODS

### Cell culture and chemicals

All cell lines were grown at 37°C in humidified incubator with 5% CO_2_ and maintained in phenol-free Dulbecco Modified Eagle Medium (Gibco) supplemented with 10% fetal bovine serum and 2mM GlutaMax (Gibco). AZD2281 (olaparib; PARPi), AZD1775 (adavosertib; WEE1i), AZD0156 (ATMi), AZD6738 (ceralasertib; ATRi), AZ’6119 (DNA-PKi), and AZ1152 (Aurora B kinase inhibitor) were synthesized internally at AstraZeneca. All compounds were dissolved in DMSO at stock concentration of 10 mM.

### Cell growth inhibition assay

SCLC NCI-H417a cells were seeded in 96-well plates and allowed to adhere overnight before being treated with DMSO, CRLX101 or the combination of CRLX101 with the different DDR inhibitors. For CRLX101 single agent assessment, treatment was for 48h before washing out the drug. DDR inhibitors were added concurrently with CRX101 and replenished after 48h. Growth inhibition (GI_50_ values) were determined at day 7 using the MTT assay in which tetrazolium MTT substrate (Sigma) was added to a final concentration of 0.25mg/mL. Following 6-8 hours incubation, formazan crystal products were solubilized in 5% SDS and 0.005M HCl (final concentrations). Optical density of each well was read at 570nM, and the absorbance readings were normalized to the DMSO control. Relative absorbance values were plotted on GraphPad Prism software to derive GI_50_ values.

### Cell line immunoblotting and antibodies

Whole cell lysates were prepared by lysing cell pellets directly in 2x Laemmli sample buffer (4% SDS, 20% glycerol, 125 mM Tris-HCl pH 6.8) and vortexing samples at highest speed for 20 seconds. Following protein concentration measurement using DC Protein Assay (Bio-Rad), protein samples were supplemented with sample reducing agents at 1x (Invitrogen) and 0.01% bromophenol blue final concentration. Samples were boiled at 95°C for 5 minutes before proteins were separated on NuPAGE 4-12% Bis-Tris protein gels (Invitrogen) by SDS-PAGE. Following protein transfer onto nitrocellulose membranes, immunoblotting analyses were performed using antibodies listed below.

| Target | Company | Catalogue No | Origin | Dilution |
| --- | --- | --- | --- | --- |
| ATM pS1981 | Abcam | ab81292 | rabbit | 1/500 |
| CDK1 pY15 | Cell Signaling Technology | 9111 | rabbit | 1/1000 |
| CHK1 pS345 | Cell Signaling Technology | 2348 | rabbit | 1/500 |
| Histone H3<br>pS10 | Cell Signaling Technology | 9701 | rabbit | 1/1000 |
| γH2AX | Cell Signaling Technology | 2577 | rabbit | 1/1000 |
| RPA32<br>pS4/S8 | Bethyl Laboratories | A300-245A | rabbit | 1/1000 |
| Vinculin | Sigma-Aldrich | V9131 | mouse | 1/5000 |

### Cell cycle analysis by flow cytometry

Cells treated with indicated compounds were harvested and fixed in ice-cold 70% ethanol overnight at −20 °C. Fixed cells were then washed with PBS before incubated in PBS containing 100 µg/ml RNase A (Thermofisher) and 50 µg/ml propidium iodide (Sigma) for 30 minutes at 37 °C. FACSCalibur (BD Biosciences) was employed to analyze samples, and data were plotted using FlowJo software.

### *In vivo* efficacy and pharmacodynamic studies

All *in vivo* studies were performed within AstraZeneca in the United Kingdom and study protocols reviewed and approved by the Home Office. All studies were performed in accordance with the Animal Scientific Procedures Act 1986 (ASPA) and AstraZeneca Global Bioethics policy. Data were reported following the Animal Research: Reporting In Vivo (ARRIVE) experiment guidelines (Kilkenny et al., 2010). For the efficacy studies, human small cell lung cancer cells (NCI-H417a) were implanted subcutaneously in athymic Fox1-nu mice. Animals were dosed with individual agents or combinations of CRLX-101 (1xwk) +/- DDR inhibitors. Animals were randomized into vehicle and treatment groups based on mean tumor volume of approximately 0.1 to 0.2 cm^3^. Animal treatment groups received either CRLX101 alone at either 4mg/kg or 5mg/kg once per week while DDR inhibitor daily dosing for olaparib was 100 mg/kg, AZD1775 120mg/kg, AZD0156 at 5mg/kg orally. Statistical significance was evaluated using a one-tailed, t test.

### *In vivo* rat studies

RccHan:WIST rats (n=7 per treatment group) were obtained from Harlan UK. The animals, aged approximately 11 weeks at the start of dosing, were allowed to acclimatize for at least 1 week and were multiple housed up to 5/cage. Water from the site drinking water supply and RM1 (E) SQC pelleted diet supplied by Special Diet Services Ltd, England was freely available. Nesting material (Tapvei Aspen Chips, Finland, Tapvei small aspen bricks and sizzle nest) and polycarbonate tunnels, Datesand were provided. CRLX-101 was formulated as a 2mg/mL PBS nanosuspension. The nanosuspension was diluted to required concentrations using PBS on the day of use. Animals dosed with CRLX-101 were given a single intravenous administration on day 1. Animals receiving olaparib, formulated in dimethyl sulphoxide diluted 1 in 10 by 10% hydroxypropyl ß-cyclodextrin in phosphate buffered saline (pH7.4), were dosed once daily by oral gavage. Where AZD2281 and CRLX-101 were dosed on the same day the olaparib (oral) dose was given approximately 1 hour after CRLX-101 (iv) dose. Topotecan was supplied by Accord Healthcare limited and formulated in 0.9% saline. Animals were dosed with topotecan daily by oral gavage. All animals were euthanased by administration of halothane.

### Flow cytometry analysis of rat bone marrow cells

Rat femurs were removed at necropsy and both ends were trimmed. Bone marrow cells were immediately flushed out with 3 ml PBS containing 50% fetal calf serum (FCS). Cell suspension was syringed and filtered through 100 μm strainer, collected by centrifugation (300 g/ 7 min/ 4°C) and washed once in HBSS containing 2% FCS and 10 mM HEPES (staining buffer). Total cell count of isolated cells was determined by automated cell counter (Countess, Invitrogen). Cell concentration was adjusted to 1×10^7^ cells/ ml in staining buffer and processed for antibody staining. CD71 and CD45 cocktail: 100 μl cell suspension was resuspended in 100 μl staining buffer containing anti-rat CD71-FITC (dilution 1:10) and CD45-PE (1:10) from Serotec. Cells were incubated with antibodies for 30 minutes RT and washed twice in staining buffer. Cell pellet stained with CD71-CD45 antibodies was resuspended in 200 μl staining buffer. This was followed by addition of 10 μl of LDS-751 cell-permeant nuclear stain (Life Technologies). CD90.1 and Lineages cocktail: 1 ml of cell suspension was resuspended in 100 μl staining buffer containing anti-rat CD90.1-APC (dilution 1:100), CD6-FITC (1:100), CD3-FITC (1:100), CD11b-FITC (1:200), Granulocytes-FITC (1:200) from BD Pharmingen and CD45RC-FITC (1:100) purchased from Serotec. After final centrifugation, 1 ml of staining buffer was added to CD90.1-Lineages-stained cells. Cells were incubated in dark for 30 minutes prior to flow cytometry analysis. Data (at least 10,000 events) were acquired on FACS AriaII (BD). Analysis was performed in FlowJo software.

### Immunohistochemistry (IHC) analysis of γH2AX in rat bone marrow and xenofraft tumor

Rat femurs or human tumour samples implanted in mice were removed at necropsy and fixed in formalin *in situ* for 48 hours prior to coring. Preserved tissue was processed into wax blocks, sectioned and stained by immunohistochemistry for the presence of γH2AX (Ventana™, Omin-UltraMap HRP, Doscovery XT Staining module; γH2AX CST 25777 @ 1:100). Sections were counter stained with haematoxylin. The slides provided for analysis were scanned using the Aperio Scanscope, converted to .TIF images (x 10 magnifications), and analysed on the KS400 image analyser.

### Haematology analysis of rat plasma samples

Blood samples to be taken from the tail vein (0.4 mL into EDTA). Haematology analysis was performed on the same day using the Siemens Advia 2120i automated haematology analyser.

## Supplemental Materials

**Figure S1:**
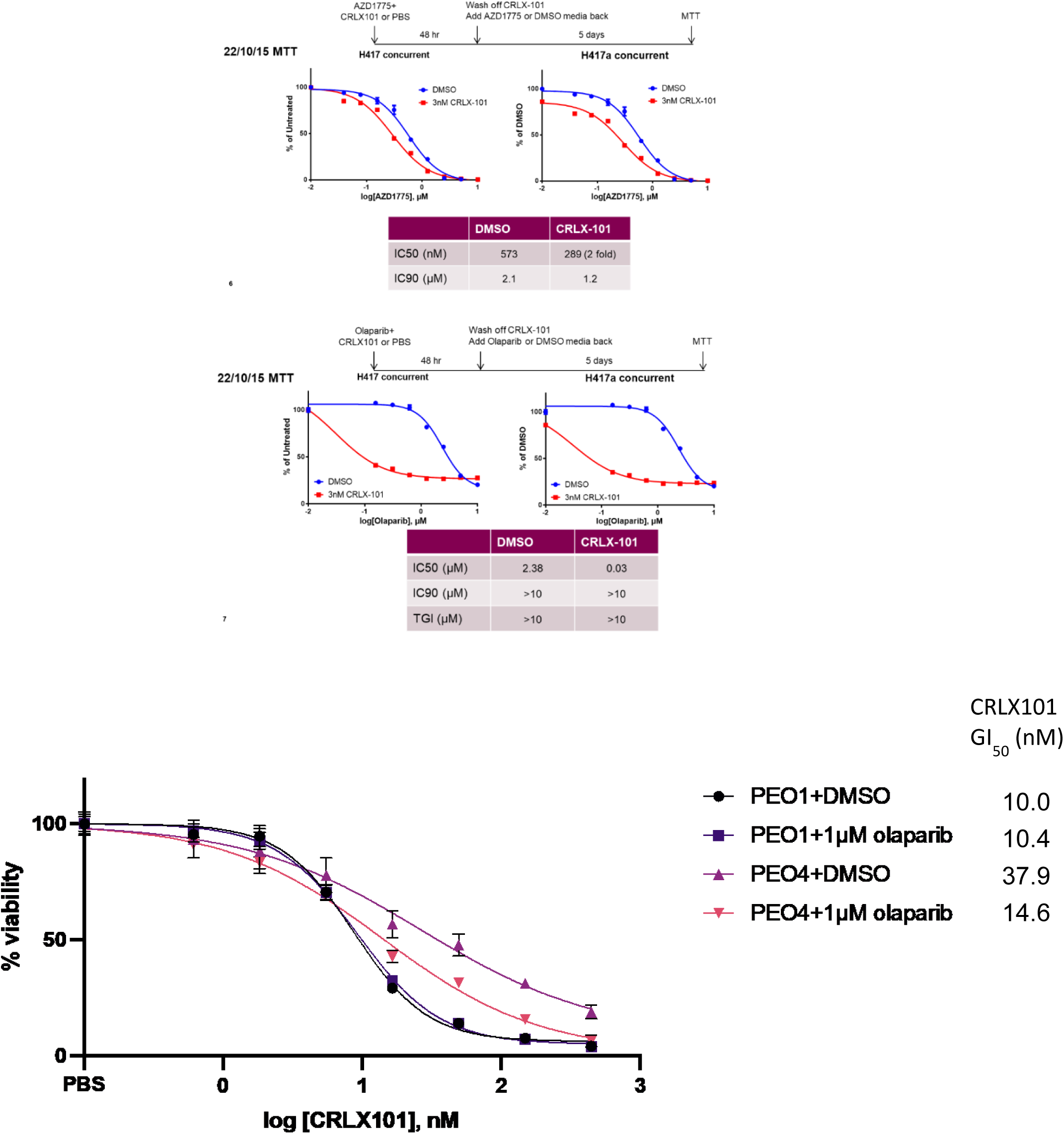
CRLX101 potentiates the efficacy of WEE1 and PARP inhibitors *in vitro*.

**Figure S2:**
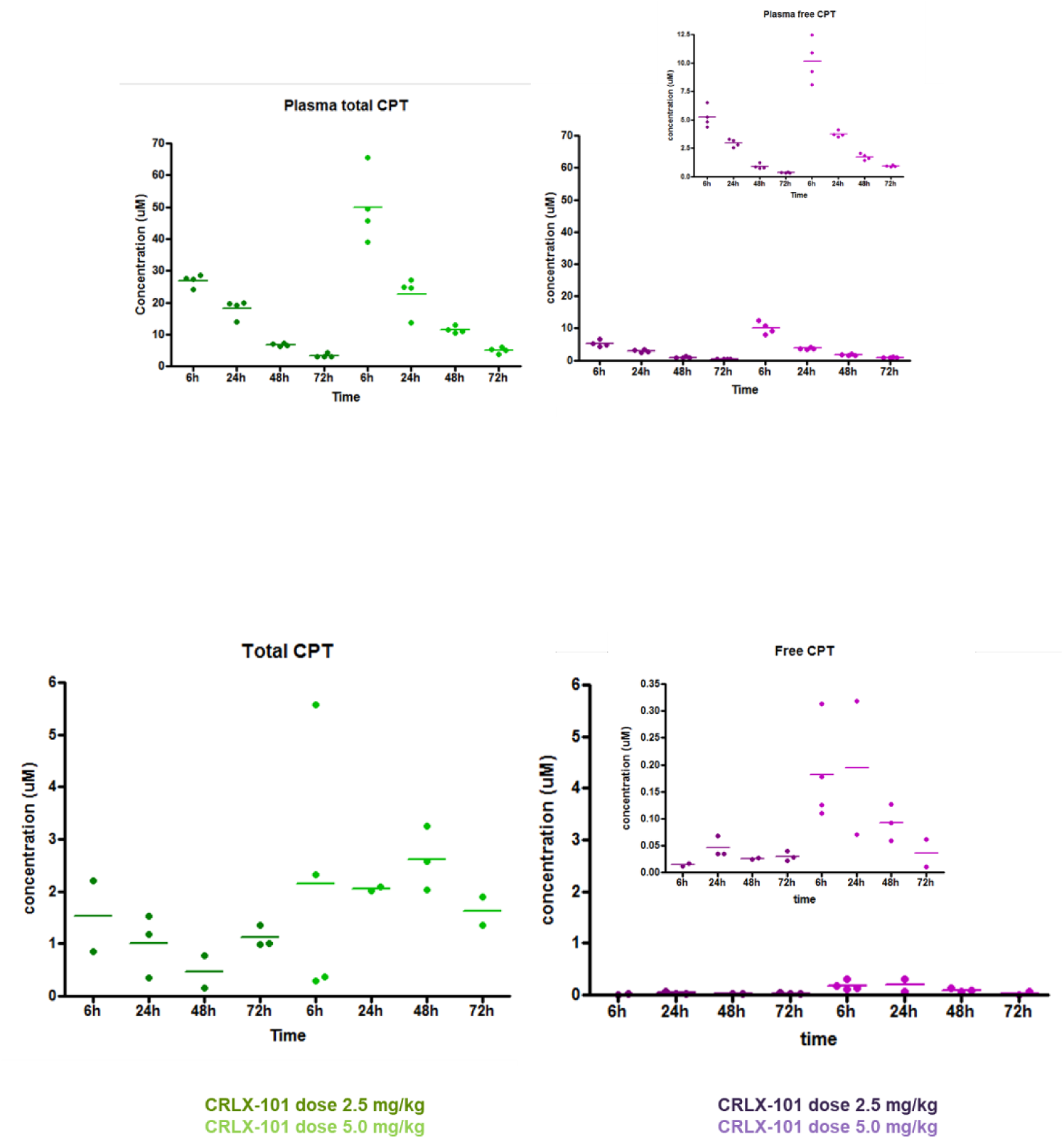
CRLX101 plasma and tumour pharmacokinetics.

**Figure S3:**
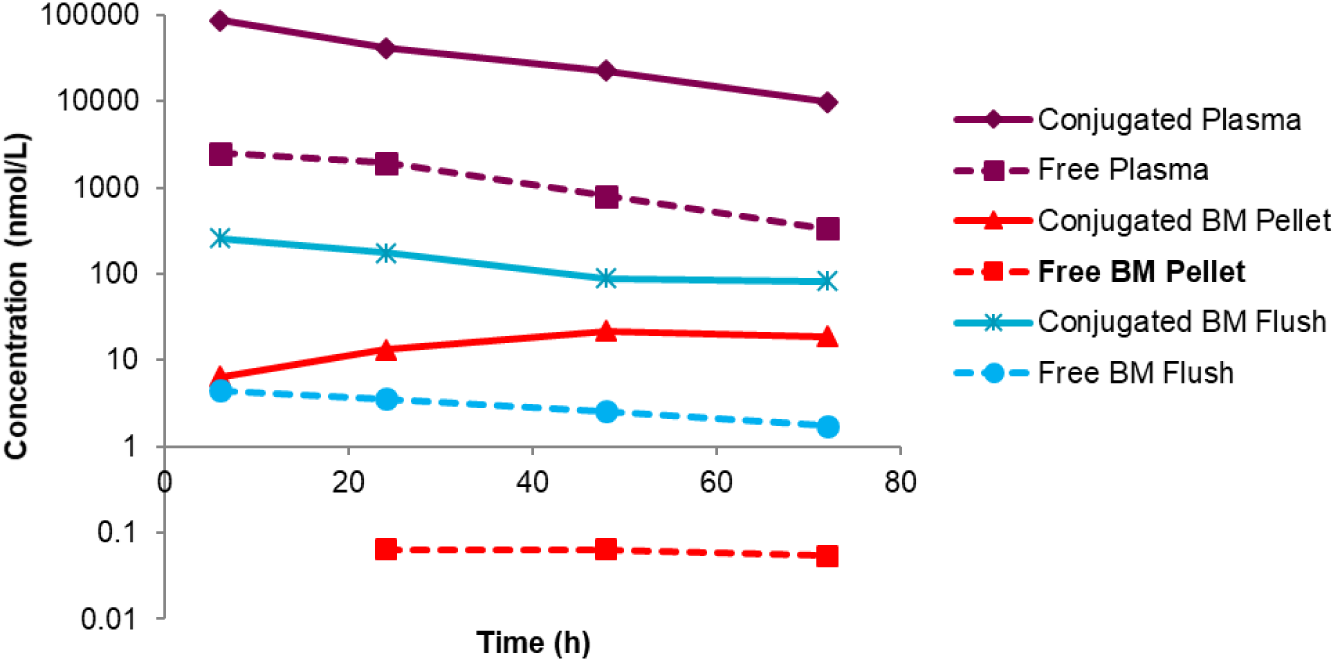
CRLX101 bone marrow pharmacokinetics.

**Figure S4:**
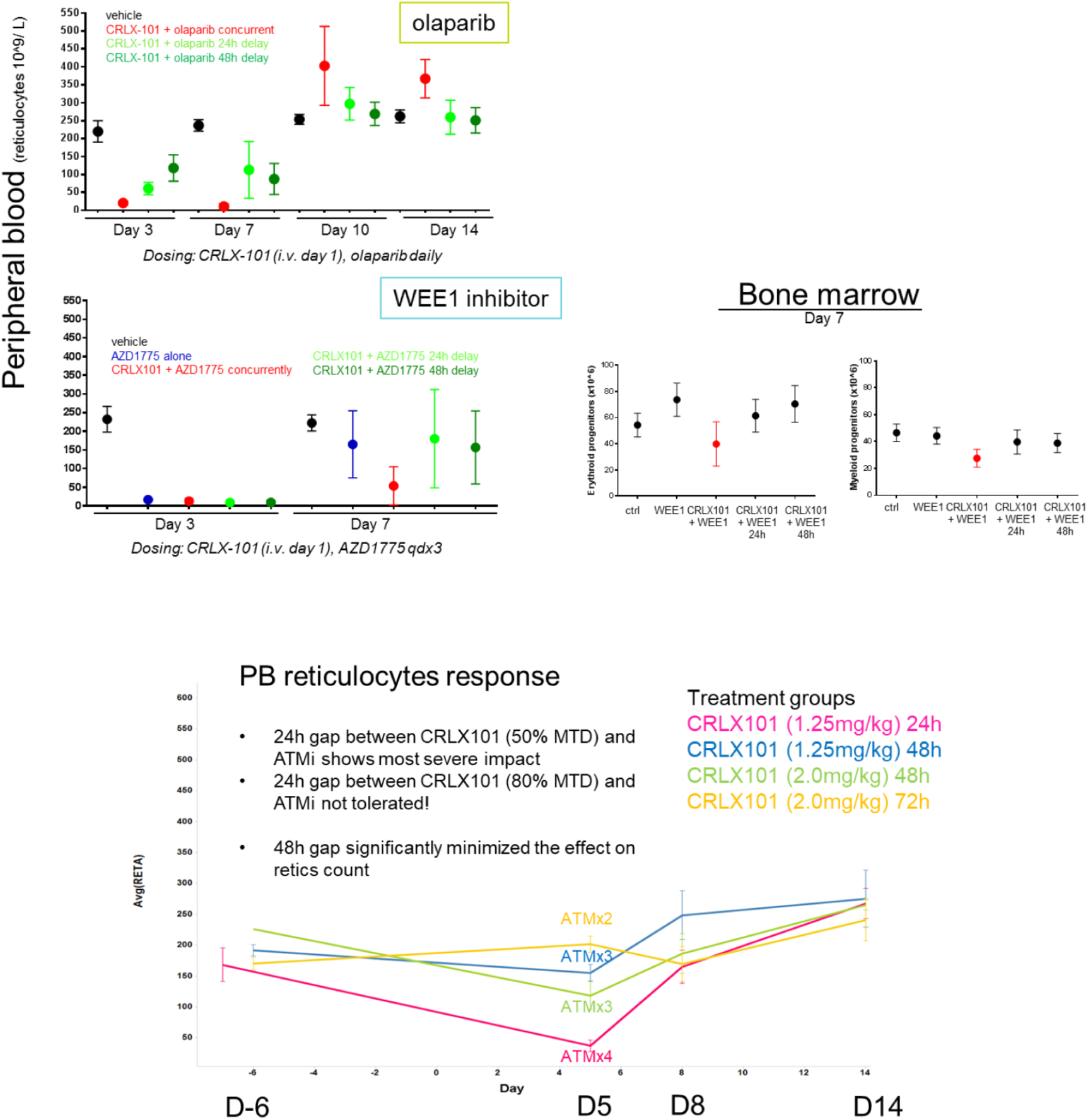
Hematological effects of CRLX101 combined with WEE1i or ATMi compared to olaparib.

**Figure S5:**
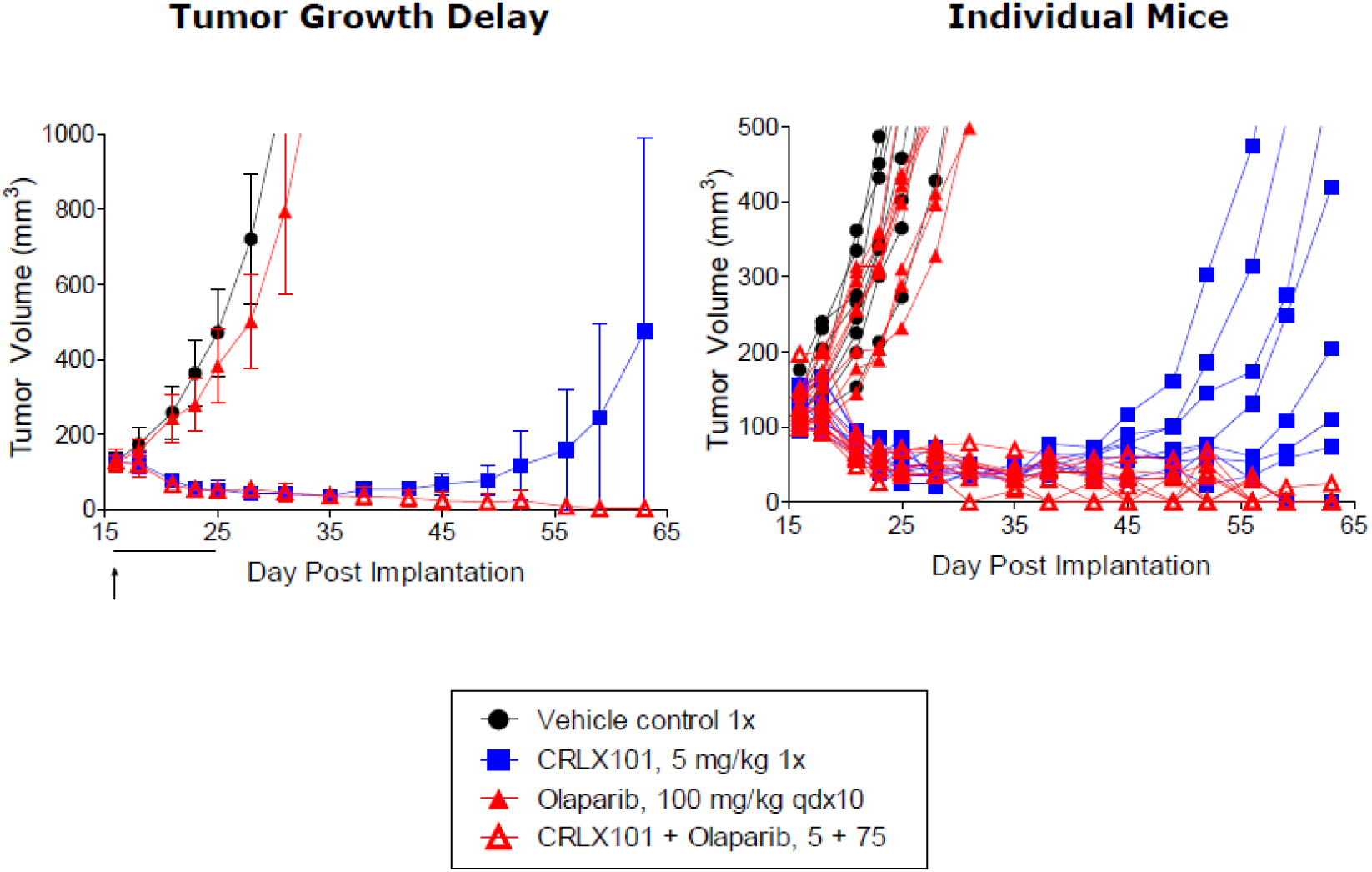
Enhanced activity of CRLX101 when in combination with olaparib in the SCLC xenograft in vivo model NCI-H480.

